# *p*Net: A toolbox for personalized functional networks modeling

**DOI:** 10.1101/2024.04.26.591367

**Authors:** Yuncong Ma, Hongming Li, Zhen Zhou, Xiaoyang Chen, Liang Ma, Erus Guray, Nicholas L. Balderston, Desmond J. Oathes, Russell T. Shinohara, Daniel H. Wolf, Ilya M. Nasrallah, Haochang Shou, Theodore D. Satterthwaite, Christos Davatzikos, Yong Fan

## Abstract

Personalized functional networks (FNs) derived from functional magnetic resonance imaging (fMRI) data are useful for characterizing individual variations in the brain functional topography associated with the brain development, aging, and disorders. To facilitate applications of the personalized FNs with enhanced reliability and reproducibility, we develop an open-source toolbox that is user-friendly, extendable, and includes rigorous quality control (QC), featuring multiple user interfaces (graphics, command line, and a step-by-step guideline) and job-scheduling for high performance computing (HPC) clusters. Particularly, the toolbox, named personalized functional network modeling (pNet), takes fMRI inputs in either volumetric or surface type, ensuring compatibility with multiple fMRI data formats, and computes personalized FNs using two distinct modeling methods: one method optimizes the functional coherence of FNs, while the other enhances their independence. Additionally, the toolbox provides HTML-based reports for QC and visualization of personalized FNs. The toolbox is developed in both MATLAB and Python platforms with a modular design to facilitate extension and modification by users familiar with either programming language. We have evaluated the toolbox on two fMRI datasets and demonstrated its effectiveness and user-friendliness with interactive and scripting examples. pNet is publicly available at https://github.com/MLDataAnalytics/pNet.

## Introduction

Functional magnetic resonance imaging (fMRI) has become an indispensable tool for exploring the brain’s function, neuroanatomy, and network dynamics. Functional connectivity (FC) is a useful measure for exploring the brain’s functional organization by quantifying covariations of different brain regions (Bastos and Schoffelen, 2015; Buckner et al., 2013; van den Heuvel and Hulshoff Pol, 2010). The emergence of large-scale fMRI datasets has provided unprecedented opportunities to characterize functional networks (FNs) and their variations across individuals, offering insights into behaviors and brain disorders (Casey et al., 2018; Harms et al., 2018; Littlejohns et al., 2020; Satterthwaite et al., 2014; Van Essen et al., 2012). Recent findings have unveiled significant individual variability in personalized FNs, underscoring the importance of delineating individual-specific FNs and their associated FC measures for investigating the brain development, aging, and neuropsychiatric disorders (Anderson et al., 2021; Cheng et al., 2023; Cohen et al., 2008; Cui et al., 2020; Cui et al., 2022; Gordon et al., 2017a; Gordon et al., 2017b; Hacker et al., 2013; Keller et al., 2024; Keller et al., 2023b; Keller et al., 2023c; Laumann et al., 2015; Li et al., 2023; Pines et al., 2022; Shanmugan et al., 2022; Wig et al., 2014; Zhou et al., 2023).

To achieve reliable and robust computation of the personalized FNs, we have developed group Sparsity-Regularized Non-negative Matrix Factorization (SR-NMF) and Group-Information-Guided Independent Component Analysis (GIG-ICA) methods that optimizes the functional coherence of FNs and enhances their independence respectively (Cui et al., 2020; Du and Fan, 2013; Li et al., 2016; Li et al., 2017). Both methods can decompose brain fMRI data into personalized FNs with spatial correspondence across individuals and have been used in a variety of brain imaging studies to capture precise individual functional network topography and their associations with personal behaviors, disease symptoms, and treatment responses (Cheng et al., 2023; Cui et al., 2020; Cui et al., 2022; Fu et al., 2023; Jing et al., 2023; Keller et al., 2023a; Keller et al., 2023c; Li and Fan, 2018, 2019; Li et al., 2016; Li et al., 2017, 2018a; Li et al., 2023; Lin et al., 2023; Pines et al., 2022; Shah et al., 2023; Shanmugan et al., 2022; Zhou et al., 2023; Zhu et al., 2021). While source code of these methods is publicly available, we recognize the need for a user-friendly and extendable toolbox with an intuitive graphical user interface (GUI) and native Python support. Additionally, the integration of quality control is essential to facilitate rigorous studies of the personalized FNs (Li et al., 2023; Zhou et al., 2023). It is worth noting that GIG-ICA has been integrated in other toolboxes developed in MATLAB platform, such as IABC (Du et al., 2023).

As part of our effort to create NiChart that enables mapping of large-scale multi-modal brain MRI data into a dimensional system of neuroimaging derived measures (https://github.com/CBICA/niCHART), we introduce personalized functional network modeling (pNet), an open-source toolbox designed for computing personalized FNs efficiently and rigorously from fMRI data. Implemented in both MATLAB and Python, pNet features a modular design, facilitating extension and customization by users to suit their individual requirements. The toolbox accepts fMRI data input in either volumetric or surface formats, compatible to fMRI data preprocessed with different preprocessing protocols, including AFNI (Cox, 1996), SPM (Penny et al., 2011), FreeSurfer (Fischl, 2012), FSL (Jenkinson et al., 2012), HCP workbench (Marcus et al., 2013), fMRIPrep (Esteban et al., 2019), and XCP (Ciric et al., 2018). pNet generates personalized FNs and provides HTML-based reports for quality control and visualization.

The rest of this paper is organized as follows. We first give an overview of pNet v1.0, including its inputs, outputs, and user interfaces. We then introduce its workflow, MATLAB GUI design, Python user interfaces, main modules, and toolbox deployment. The effectiveness and user-friendliness of our toolbox are demonstrated using two fMRI datasets. We demonstrated pNet’s features in terms of compatibility, flexibility, reliability, intuitive visualization, and reproducibility. These features have the potential to enhance user experiences and further benefit studies of subject-specific brain’s functional topography.

## Methods

### Toolbox overview

pNet v1.0 is an open-source toolbox with a modular design for computing personalized FNs. As illustrated in Figure 1, pNet consists of extendable modules of input, computation and output, as well as multiple user interfaces. It supports fMRI input in either volume or surface formats and can load existing group-level FNs as an optional input to derive personalized FNs. Implemented with native code in both MATLAB and Python platforms, pNet offers two different methods, SR-NMR and GIG-ICA that optimizes the functional coherence of FNs and enhances their independence respectively (Cui et al., 2020; Du and Fan, 2013; Li et al., 2016; Li et al., 2017), to compute personalized FNs. It provides user interfaces tailored for different user preferences and computation environments, including a MATLAB-based GUI, interactive step-by-step command line guides, and scripts for workstation and cluster computation. pNet outputs group-level and personalized FN results along with preconfigured visualizations, quality control results, and HTML-based computation reports. pNet is publicly available at https://github.com/MLDataAnalytics/pNet.

**Figure 1.**
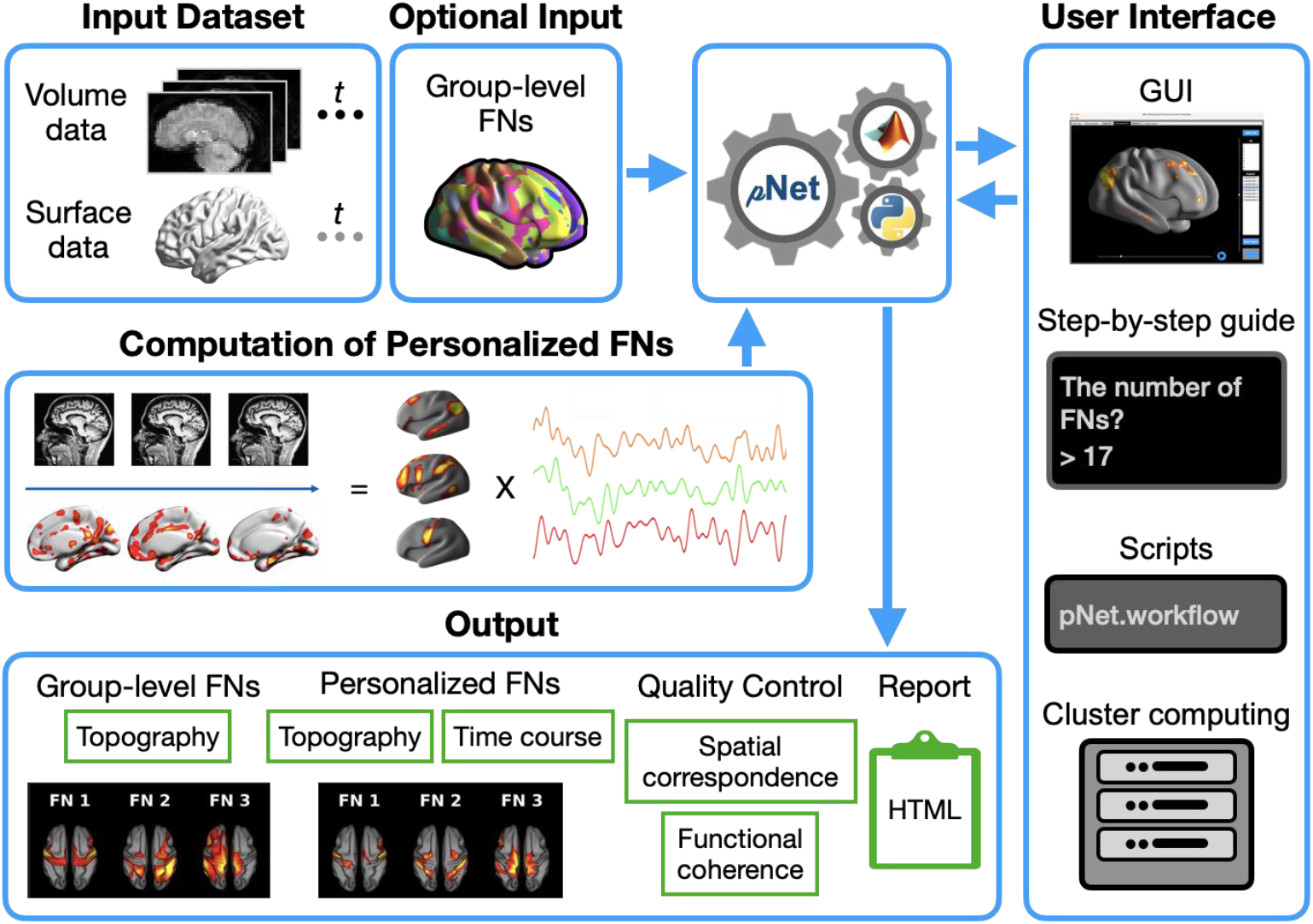
Illustration of pNet’s workflow. pNet accepts fMRI data in either volumetric or surface formats as input. Optionally, group-level FNs can be used as an additional input to guide the computation of personalized FNs. pNet offers several user interfaces, including a GUI, a command-line based interactive step-by-step guide, and a variety of scripts for running the toolbox on workstations and HPC clusters. It computes and outputs personalized FNs and provides preconfigured visualizations of FNs and quality control results in HTML format. pNet works on both MATLAB and Python platforms.

### GUI Design

The MATLAB GUI is designed to align with the pNet workflow, as illustrated in Figure 2. It comprises 6 tabs dedicated to setting input data, adjusting parameters for computing personalized FNs, visualizing computational results, and checking quality control results (Figure 2A). Interactive visualization options within the GUI allow users to visually examine multiple FNs, change views, and compare FNs obtained from different scans (Figure 2B and C). In addition, users can check the binarized functional atlas derived from group-level or personalized FNs (Figure 2D). The visualization of statistical results for personalized FNs also features interactive and synchronized display to ensure a consistent user experience.

**Figure 2.**
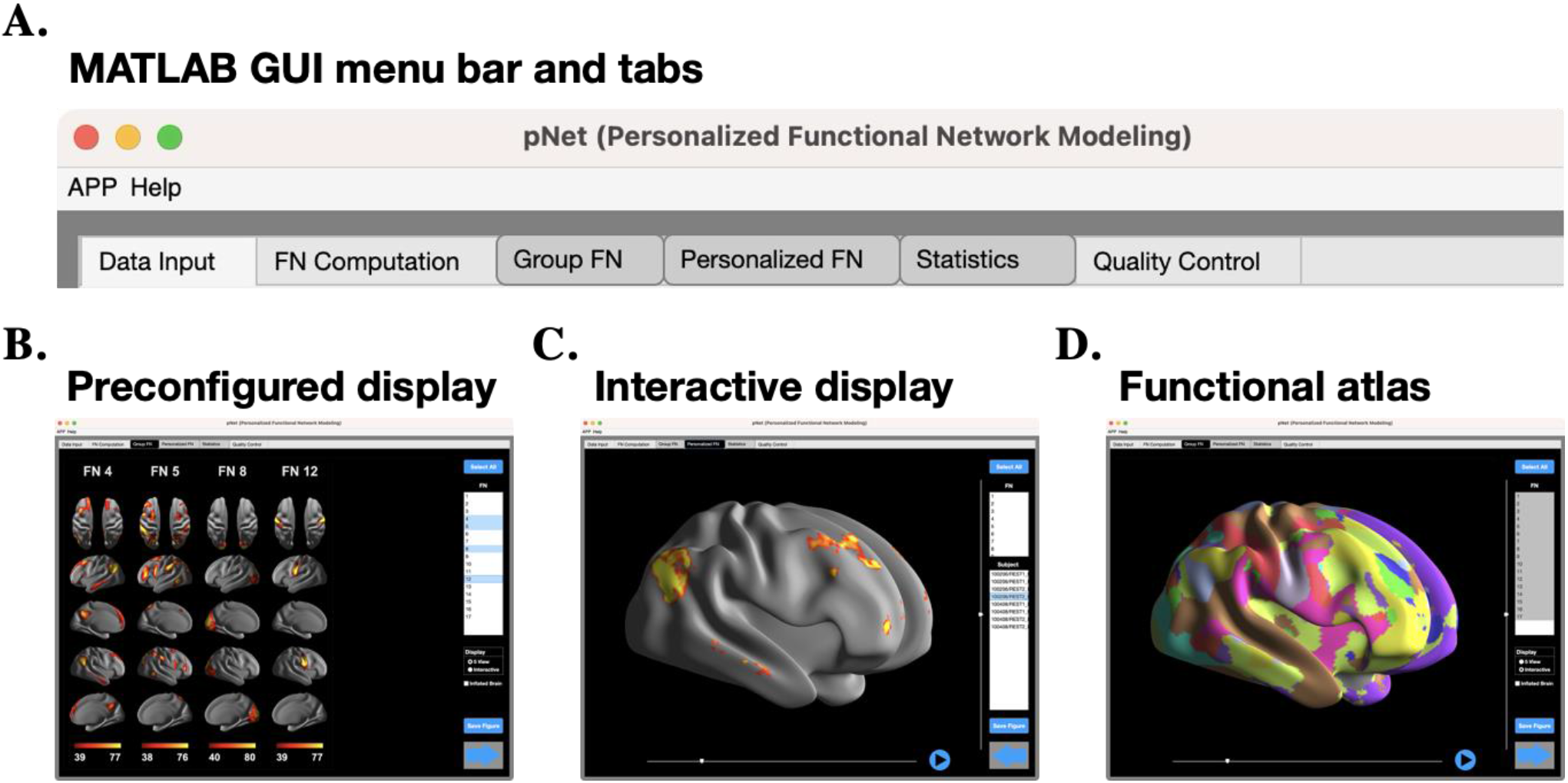
GUI design. (A) The menu bar and six tabs on the MATLAB GUI; (B) Preconfigured displays of four selected group-level FNs computed based on the HCP dataset; (C) Interactive display of a single personalized FN; and (D) Display of a binarized atlas created from the group FNs. For all intensity maps, spatial weights of each FN are displayed with the values ranging from the 50th to the 99.8th percentile.

### Python user interfaces

The Python version of pNet offers a variety of user interfaces (Figure 3). First, a step-by-step guide prompts users with questions and choices to configure a suitable workflow. This process automatically generates a Python script with detailed descriptions for the computation settings. Second, several example scripts are provided to setup a workflow with either minimal parameters or advanced settings for creating brain templates and customized the computational parameters of personalized FNs. For the computation on HPC clusters, users can configure the cluster environment with advanced settings for computation resources including CPU thread and memory allowance. In addition, pNet provides open-source internal functions, independent to the overall workflow, for users to take the advantage of the built-in FN models and preconfigured visualizations.

**Figure 3.**
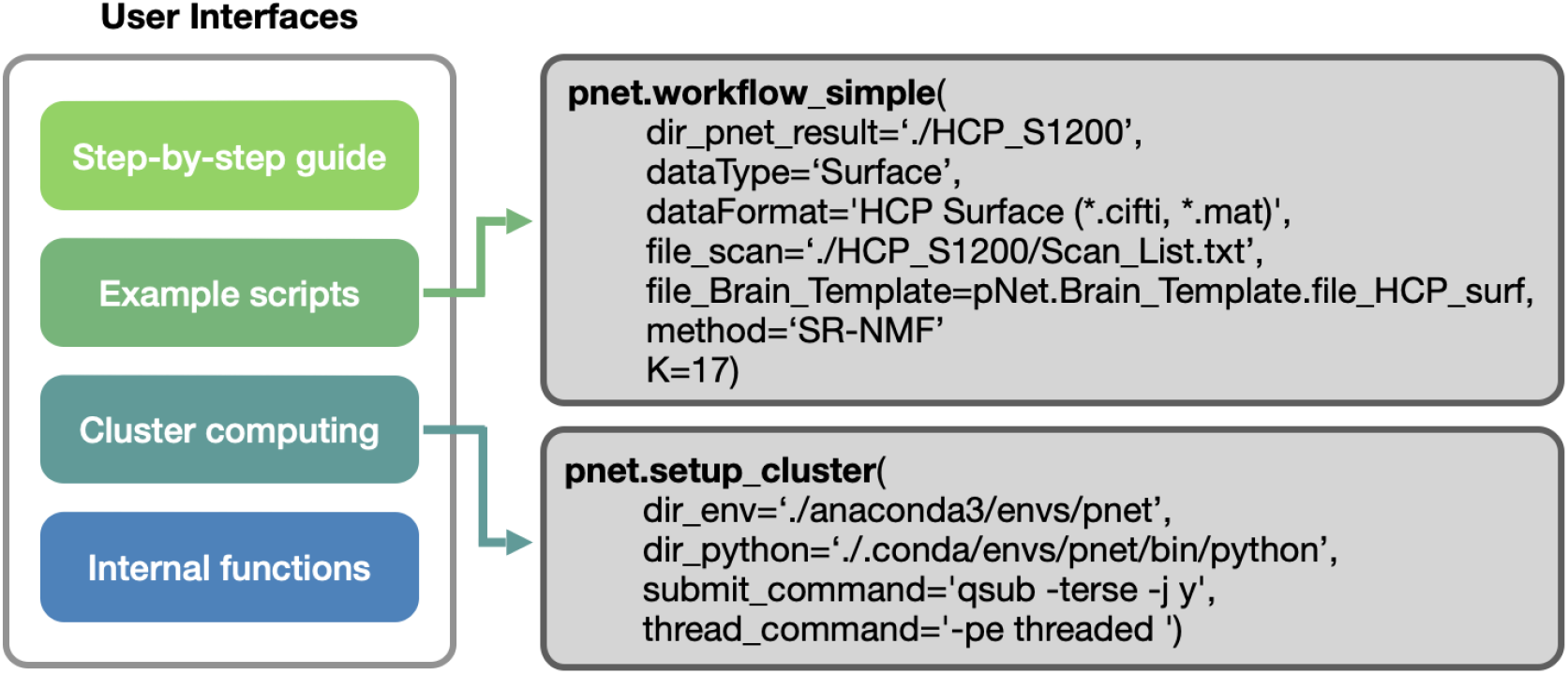
User interfaces of the Python version. The Python version offers different user interfaces, with illustrations of minimal setup options for fast deployment on workstations (pNet.workflow_simple) and advanced settings for HPC clusters (pNet.setup_cluster), respectively.

### Input module

The input module is designed to load preprocessed fMRI scans and their corresponding brain template files. It supports both volumetric and surface fMRI data stored in a variety of file formats and allows flexible file searching and modification. For volumetric data, both NIFTI and MAT formats are supported, which stores 3D spatial + 1D temporal fMRI signals. The surface data typically comprises a set of gray matter vertices, each attached with an 1D temporal fMRI signal. pNet supports four surface data formats, including CIFTI, MGZ, MGH, and MAT. Specifically, CIFTI, MGZ, and MAT formats store data for each scan in a single file, and the MGH format employs two files to store data for the left and right hemispheres separately. For the MGH format, pNet merges the directories of the two files into one. In addition, pNet supports a combination of surface and volumetric data types stored in HCP grayordinate format.

pNet obtains its input by utilizing a text file that contains full directory information for the fMRI scans. Additionally, it allows users to curate customized fMRI datasets through an intuitive GUI for searching fMRI scans by file names, extensions, or manually select multiple scans. pNet combines the search results to generate a dataset file for subsequent computation of the FNs. Furthermore, it allows manual modifications of the scan list file to further customize the fMRI dataset.

Since fMRI signals may exhibit a limited signal-to-noise ratio (SNR), multiple scans are often acquired for the same individual to enhance fMRI data quality. In response to this need, pNet offers flexibility for concatenating multiple fMRI scans to form a single continuous one. When concatenating data, pNet can automatically use the sub-folder name (where the fMRI files are stored) as the subject ID. This simplifies the process by associating each concatenated scan with a specific subject. Alternatively, users can load text files containing subject IDs. These files should follow the same order as the corresponding fMRI files. This approach ensures accurate alignment between subject IDs and their respective scans. The segments of the same fMRI scan can be stored in sub-folders with a same subject ID and the file information of segments can also be stored in text files. When data concatenation is enabled, a single set of personalized FNs will be generated for each subject; otherwise, each fMRI scan segment will have its own set of personalized FNs.

### FN computation module

This module is designed to setup and carry out the computation of personalized FNs. In particular, pNet computes the personalized FNs with either SR-NMF or GIG-ICA, both of them yielding personalized FNs that are regularized with group-level FNs for establishing spatial correspondence across individuals (Cui et al., 2020; Du and Fan, 2013; Li et al., 2016; Li et al., 2017).

For GIG-ICA, any group ICA method can be used to compute group-level FNs for guiding the computation of the personalized FNs, such as GIFT and MELODIC (Beckmann and Smith, 2004; Correa et al., 2005), and then the group-level FNs can be used as guidance information for computing the personalized FNs (Du and Fan, 2013). pNet provides scripts for computing group-level FNs using GIFT (https://trendscenter.org/software/gift/).

For SR-NMF, the personalized FNs are computed jointly for a group of subjects regularized by group sparsity (Li et al., 2016; Li et al., 2017). Since such a joint computation of personalized FNs does not scale well to large-scale datasets due to its prohibitive computational memory consumption (fMRI data of all scans has be considered simultaneously), the similar strategy of GIG-ICA is adopted to compute the personalized FNs with regularization information provided by group-level FNs, in conjunction with a bootstrapping strategy by sampling the large datasets (Cui et al., 2020).

To further enhance reliability, a spatial correspondence constrain is integrated into GIG-ICA and SR-NMF to ensure one-to-one correspondence between personalized FNs and their group-level counterparts. Hence, the spatial correspondence is guaranteed for group or individual level comparisons.

### Output module

pNet outputs personalized FNs along with QC and visualization results, which can be accessed through the GUI tabs (Figure 2A). Moreover, computational log files are generated to store information of the fMRI dataset and parameters used for the computation, as well as intermediate results in folders named as “Data_Input”, “FN_Computation”, “Group_FN”, and “Personalized_FN”, respectively. FNs and their corresponding time courses are stored in separate files, and based on the time courses, functional connectivity (FC) measures can be computed as Pearson correlation values (Zhou et al., 2023). QC results are stored in a folder named “Quality_Control”. An HTML file named “Report.HTML” provides links to HTML-based QC reports of individual scans or subjects in their corresponding subfolders within folder “Personalized_FN”.

### Quality control

Quality control is implemented to ensure that personalized FNs have higher functional homogeneity than their group-level counterparts and maintain good spatial correspondence with their group-level counterparts (Li et al., 2023; Zhou et al., 2023). Particularly, the functional coherence of each FN is measured by a weighted mean of the correlation coefficients between the time courses of all the voxels within the FN and its centroid time course, which is calculated as a weighted mean time course over the FN with its voxel-wise loadings as weights. The spatial correspondence between each personalized FN and its group-level FN is measured based on their pairwise spatial correlation coefficients. Specifically, each personalized FN is deemed to maintain correspondence with its corresponding group-level counterpart if its spatial correspondence value is greater than all pairwise spatial correlation coefficients between the personalized FN under consideration and all other group-level FNs. The quality control results are generated with the computation of FNs.

### Statistical analysis

pNet incorporates four commonly-used statistical tests for analyzing the personalized FNs. Particularly, one-sample t-test and Wilcoxon signed rank test are integrated to assess the significance of personalized FNs (voxels for volumetric data and vertices for surface data) across subjects. Two-sample t-test and Wilcoxon signed rank test are included to facilitate group comparison of the FNs. Covariates, such as demographic data, can be included in the statistical analyses. The resulting statistical results, including spatial maps of p or t/z values, can be visualized with the GUI. To mitigate the risk of type I errors, p-values can be further adjusted using the false discovery rate (FDR) correction method (Lieberman et al., 2009).

### Toolbox Development

pNet is developed with native MATLAB and Python code respectively, facilitating code integration with other packages and toolboxes available on the two platforms. As an open-source software, pNet is available on GitHub (https://github.com/MLDataAnalytics/pNet), promoting collaborative development and providing a platform for receiving valuable feedback for enhancement and future development. The MATLAB version has been tested on macOS, Linux, and Windows operating systems. It provides several execution methods, including its GUI design, which can function as a MATLAB GUI or be installed as a MATLAB APP. The Python version is tested on Anaconda (https://www.anaconda.com). It can also be built into a Docker container (https://www.docker.com) to facilitate easy adoption of the toolbox. Moreover, scripts are provided to facilitate job-scheduling for computing on HPC clusters. Our scripts have been tested on the CUBIC (a RedHat Enterprise Linux-based HPC cluster, https://www.med.upenn.edu/cbica/cubic.HTML) at the University of Pennsylvania.

### Expandability

With the rapid evolution of the fMRI field, new data formats and modeling methods are being introduced at a rapid pace. To facilitate expandability, pNet is developed with a modular design, allowing for addition of new functions and methods. Additionally, pNet encourages future software development to include new data formats and personalized functional network models, and statistic methods. With the availability of both MATLAB and Python versions, pNet allows users with different coding preferences for further expansion.

#### Toolbox Validation

To evaluate the effectiveness and versatility of pNet, we applied it to two different resting state fMRI (rsfMRI) datasets with fMRI scans preprocessed in their specific studies. In the first application, we tested pNet on volumetric fMRI data from the UK Biobank dataset (Littlejohns et al., 2020). We loaded two sets of precomputed group-level FNs (k=17 and 21) from our existing multi-scale FN study (Zhou et al., 2023) and another study (Miller et al., 2016) for testing the MATLAB version. For this dataset, we chose a small subset (10 subjects) to calculate personalized FNs, thereby demonstrating the ease-of-use of the GUI-based pNet and convenience for visual examinations and comparisons. For the second application, we tested pNet on surface fMRI data of 478 subjects from the Human Connectome Project (HCP) dataset (Van Essen et al., 2012). We used the Python version to obtain both group-level and personalized FNs, as well as the quality control results, demonstrating its usability in an HPC cluster environment. The results of this toolbox validation confirmed its compatibility, flexibility, reliability, and efficiency, thus establishing it as an asset for exploring brain functional networks and their relationship to behavior or mental disorders.

The toolbox comes with two sets of precomputed group-level FNs (k=17), computed with SR-NMF based on the HCP young adult dataset and a subset of UKBB from the iSTAGING study (Zhou et al., 2023), respectively. Specifically, the HCP derived group-level FNs are in surface format, and the UKBB derived ones are in volumetric format. The group-level FNs can be loaded to compute personalized FNs on other datasets. This eliminates the computationally heavy burden of computing group-level FNs and ensure reliable comparisons across datasets as well as reproducibility.

### Application 1: Utilizing precomputed group-level FNs to obtain personalized FNs

We adopted a subset of the UK Biobank (UKBB) dataset, which passed quality control in our previous study (1571 subjects, 695 males, ages 45-79, 420 volumes per scan) (Zhou et al., 2023). A standardized preprocessing pipeline was used (Alfaro-Almagro et al., 2018). Subsequently, preprocessed data were transformed into the MNI space and stored in NIFTI format. In this application, we employed two sets of precomputed group-level FNs, including 17 FNs from our previous study (Zhou et al., 2023) that were derived from a multi-cohort iSTAGING dataset (4186 subjects, ages 22-97), and 21 group-ICA components obtained from UKBB dataset (Miller et al., 2016). These results were loaded from the GUI, and subsequently used to compute personalized FNs for a subset of the UKBB dataset (10 subjects, 490 time points per scan). SR-NMF and GIG-ICA were used for computing personalized FNs guided with the two sets of group-level FNs, respectively.

Figure 4B and D show results of personalized FNs derived via SR-NMF and GIG-ICA, respectively. The GUI can display preconfigured visualizations (Figure 4A and C) and also allows for interactive visualization of results (Figure 4B and D). Synchronized display is also available to facilitate swift comparisons between group-level and personalized FNs with customizable display settings. For example, users can compare group-level and personalized FNs with the same FN selection (Figure 4A and B) and view center point (Figure 4C and D).

**Figure 4.**
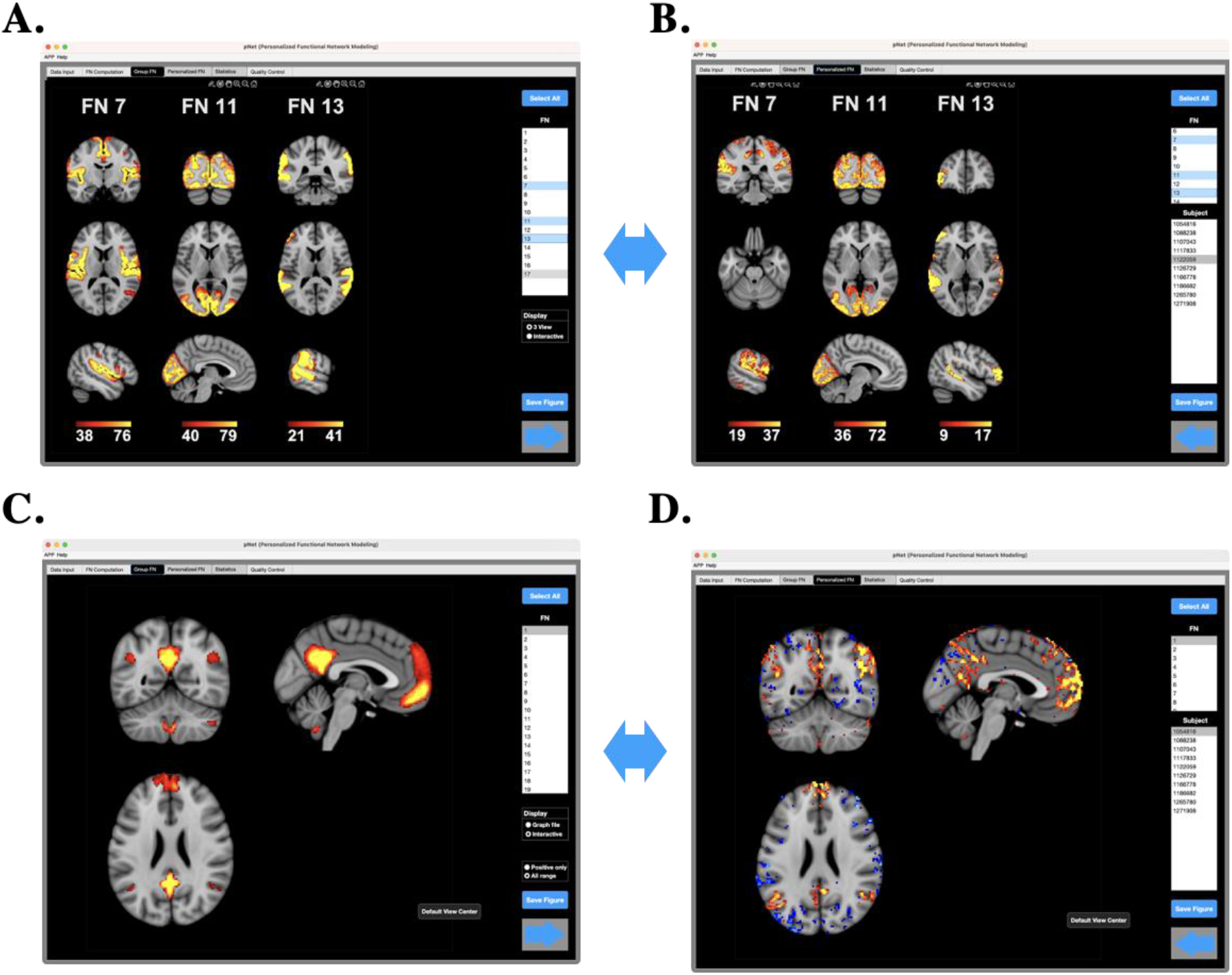
Visualization of group-level and personalized FNs, including preconfigured visualization of group-level (A) and personalized (B) FNs for K=17, as well as interactive displays of a single group-level (C) and corresponding personalized (D) FN for K=21. Double-sided blue arrows denote synchronized displays between group-level and personalized FNs.

### Application 2: HCP dataset

We utilized a subset of minimally preprocessed resting-state fMRI (rsfMRI) data from the HCP S1200 dataset, consisting of 478 subjects aged between 22 and 35 years. Each subject underwent two or four 15-minute scans (two resting-state fMRI sessions with two scans in each session, yielding 1200 volumes per scan). Details of this minimal preprocessing pipeline can be found in (Glasser et al., 2013). We selected surface-based data in CIFTI format for evaluating pNet. A built-in brain surface template is used to provide spatial information for visualizations. The fMRI signals were further processed using XCP-D (https://xcp-d.readthedocs.io/en/latest/index.HTML, FWHM = 5mm, band pass filtering at 0.01-0.1Hz, 36 parameters for nuisance regression). We computed personalized FNs for each scan separately with default parameter settings on an HPC cluster. To enhance the computation efficiency, the whole process was automatically divided into multiple bash jobs to take the advantage of parallel computation in a cluster environment.

Figure 5 shows part of the results generated by pNet, including group-level FNs and personalized FNs of a randomly selected fMRI scan (Figure 5 A and B). Figure 5C shows QC results in terms of spatial correspondence of the personalized FNs, reflecting the spatial similarity between personalized FNs and their group-level counterparts higher than those between unmatched FNs, i.e., each personalized FN had higher spatial similarity with the corresponding group-level FN than with others and the difference (delta spatial correspondence) was significantly larger than 0. The functional coherence of the personalized FNs was significantly higher than their corresponding group-level FNs’ (p value < 0.0001, via Wilcoxon signed-rank test) (Figure 5D).

**Figure 5.**
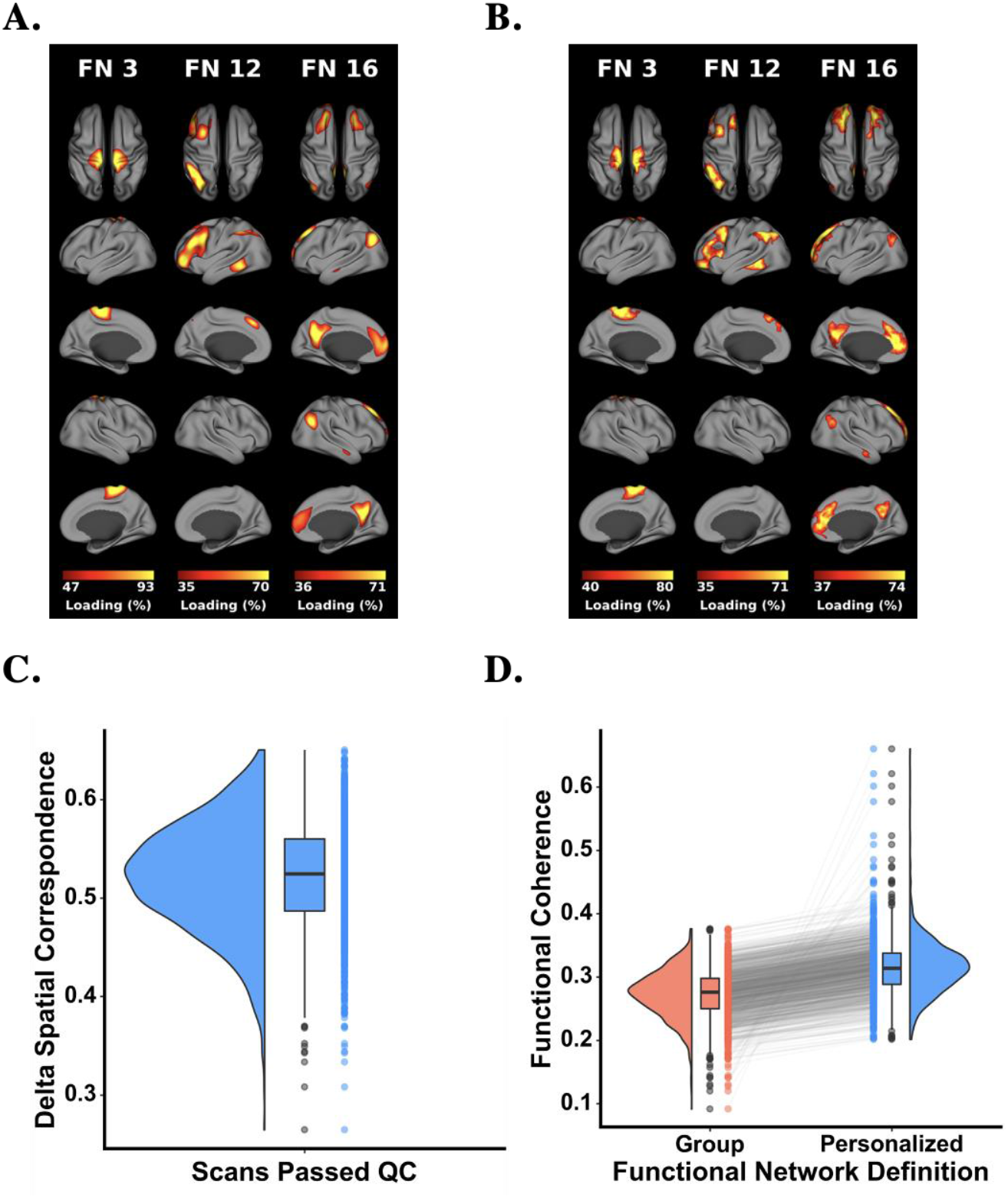
Illutration of results of the HCP dataset with 17 FNs, including three select group-level FNs (A), their corresponding personalized FNs obtained from a single fMRI scan (B), as well as QC results of spatial correspondence (C) and average functional coherence of group-level and personalized FNs of individual fMRI scans.

pNet also outputs a computation report in HTML format to facilitate visual examination of the computational results. As illustrated in Figure 6, the report includes a brief description about the dataset, main settings of the workflow, visualization results of both group-level and personalized FNs, and QC results. For fast web page navigation, visualization results of FNs are stored at a lower resolution and the report provides visualization of randomly selected personalized FNs while all personalized FNs can be visually checked through hyperlinks to individual reports.

**Figure 6.**
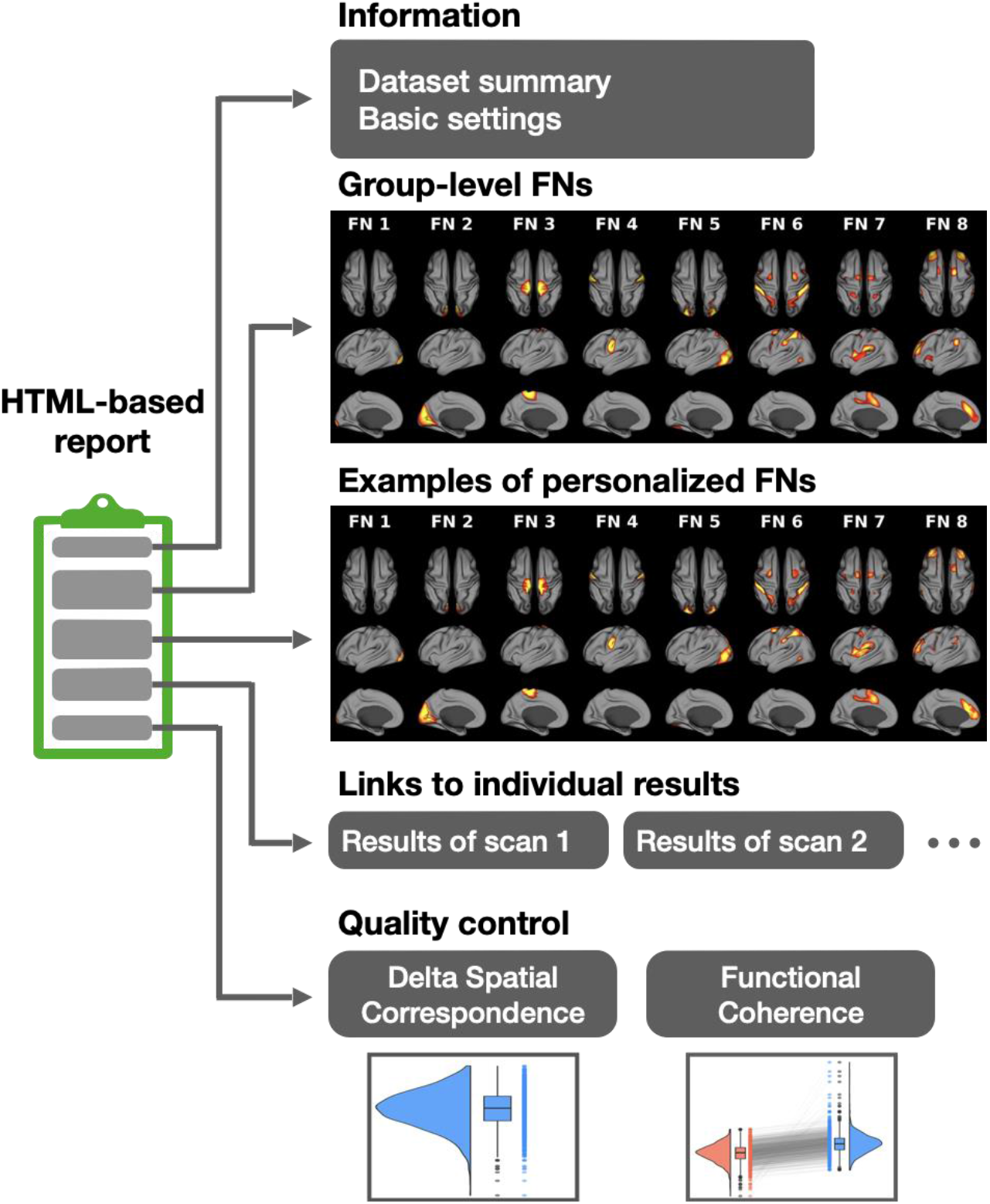
HTML-based computation report. pNet generates an HTML-based report for users to navigate through the results. It consists of basic information of the dataset and workflow, visualization results of group-level FNs, and examples of personalized FNs. All personalized FNs are accessible via hyperlinks. Results of quality control are also included in the report.

## Discussion

We introduced pNet v1.0, a user-friendly toolbox designed for computing personalized FNs from fMRI data. This toolbox encapsulates two personalized functional network modeling methods, SR-NMF and GIG-ICA, with native source code in both MATLAB and Python. pNet v1.0 is designed primarily to assist neuroscientists, psychiatrists, and researchers, with interactive visualizations. pNet offers multiple user interfaces, including a GUI, an interactive command-line based step-by-step guide, and scripts for computation on both personal computers and high-performance clusters. In addition, the modular design of pNet v1.0 allows for accommodating future expansions for other fMRI data formats, brain decomposition and statistical methods, ensuring its continued development and expandability. We have demonstrated its effectiveness and user-friendliness by testing it on two fMRI datasets.

### Comparisons with other similar toolboxes

Several widely used toolboxes offer similar functions for computing functional networks from fMRI data, such as GIFT, MELODIC, and HINT (Beckmann and Smith, 2004; Correa et al., 2005; Lukemire et al., 2020). These toolboxes follow a similar workflow that encompasses the loading of an fMRI dataset, setting up the computation parameters, generating FN results, and providing visualizations through a GUI. Despite their methodological differences, we briefly compare them to our toolbox regarding data format support, visualization options, and quality control (Table 1). With the increasing diversity in the fMRI field, a variety of data formats have been introduced to facilitate data analyses. Surface-based data have been developed to significantly reduce data size and benefit from precise surface-based image registration (Glasser et al., 2013). Our toolbox supports CIFTI, MGH, MGZ, and the MATLAB based formats. pNet also allows customizable data organization to support multi-cohort datasets. Users can easily modify the file selection in the GUI to curate the desired dataset, allowing for the removal of unwanted scans or those of insufficient quality.

**TABLE 1.** Comparison of the main features of the available toolboxes.

|  | *GIFT* | *MELODIC* | *HINT* | *pNet* |
| --- | --- | --- | --- | --- |
| *Interface* | GUI, command | GUI, command | GUI, command | GUI, command |
| *Standalone software* | × | ✓ | × | ✓ |
| *Volumetric format* | 2 | 2 | 1 | 2 |
| *Surface format* | × | 2 | × | 4 |
| *Preconfigured visualization* | × | × | × | ✓ |
| *Synchronized display* | × | × | × | ✓ |
| *Quality Control* | × | × | × | ✓ |
| *HTML-based report* | × | × | × | ✓ |

Recognizing the growing need for both automatic and customizable visualizations of FNs, pNet offers two unique features in addition to common interactive display in GUI. Firstly, it includes preconfigured visualization figures saved in the result folder, allowing users to visually examine group-level and personalized FNs in GUI, HTML-based reports and result folders. Secondly, it features a synchronized display for swift comparisons between group-level and personalized FNs. Uniquely, our toolbox integrates quality control directly into brain decomposition methods, and provides an additional quality control in the report. Users can also examine the quality assurance indices in the quality control for comprehensive information. In addition, pNet provides HTML-based reports to navigate through the results without using the toolbox, enhancing user-friendliness and reproducibility.

### Compatibility and computational requirement

pNet is available on both MATLAB and Python platforms. The MATLAB version requires MATLAB no older than R2021A and can run on macOS, Linux, and Windows systems, as well as Linux-based clusters. pNet also operates as a standalone software with the freely available MATLAB Runtime. All GUI version experiments were conducted on a 2019 Mac Pro, equipped with a 2.7GHz 24-core Intel Xeon W-3265M CPU and 256GB memory. The compatibility of Linux and Windows systems was primarily tested using Parallel Virtual Machine, and the cluster-based scripts was tested on the RedHat Linux-based cluster (CUBIC) at the University of Pennsylvania. The Python version was tested with anaconda (Python 3.8), which ensures a reproducible computation environment across different operation systems.

pNet is designed to run on personal computers with limited computational resources, high-end workstations and HPC clusters. pNet uses a two-stage strategy for the computation of personalized FNs, which first computes or loads group-level FNs, then computes personalized FNs with group-level results as initialization. The group-level FN computation typically requires a large amount of memory space to load numerous fMRI scans. For instance, loading 100 resting-state fMRI scans from the HCP S1200 dataset to compute FNs requires around 100GB of memory in total. In this case, we recommend allocating 100GB of memory for each bootstrap repetition. For computing the personalized FNs, fMRI data of single subjects will be loaded into memory each time. We recommend a memory requirement set as twice as the data size, plus the need to load the computation software (0.5-1GB). pNet also supports parallel computation, which typically requires a substantial amount of memory. The toolbox includes automatic hardware configuration detection in GUI mode and provides a rough memory usage estimation based on the CPU core number.

### Limitations and future works

Currently, pNet supports two personalized FN modeling methods. We will integrate more advanced brain decomposition methods, such as deep learning-based models (Li et al., 2023; Li et al., 2018b), to further enhance the toolbox’s capability. Currently, our Python version uses PyTorch (https://pytorch.org) to accelerate matrix computation with CPU, which can be further improved with GPU-based acceleration in term of the computational efficiency.

## Conclusions

pNet offers user-friendly functions for computing personalized FNs from fMRI data. This toolbox delivers an integrated solution for computation, visualization, quality control, statistical analyses, significantly simplifying and enhancing research in this domain.

## Acknowledgments

This work was supported in part by NIH grants of U24NS130411, R01EB022573, R01AG066650, and R01MH120811.

